# Reversal of the adipostat control of torpor during migration in hummingbirds

**DOI:** 10.1101/2021.05.13.443997

**Authors:** Erich R. Eberts, Christopher G. Guglielmo, Kenneth C. Welch

**Affiliations:** Department of Biological Sciences, University of Toronto Scarborough, 1265 Military Trail, Toronto, Ontario, Canada, M1C 1A4; Department of Ecological and Evolutionary Biology, University of Toronto, 25 Willcocks Street, Toronto, Ontario, Canada, M5S 3B2; Department of Biological Sciences, Advanced Facility for Avian Research, University of Western Ontario, 1393 Western Road, London, Ontario, Canada, N6G 1G9

## Abstract

Many small endotherms use torpor to reduce metabolic rate and manage daily energy balance. However, the physiological “rules” that govern torpor use are unclear. We tracked torpor use and body composition in ruby-throated hummingbirds (*Archilochus colubris*), a long-distance migrant, throughout the summer using respirometry and quantitative magnetic resonance. During the mid-summer, birds entered torpor at consistently low fat stores (∼5% of body mass), and torpor duration was negatively related to evening fat load. Remarkably, this energy-emergency strategy was abandoned in the late summer when birds accumulated fat for migration. Migrating birds were more likely to enter torpor on nights when they had higher fat stores, and fat gain was positively correlated with the amount of torpor used. These findings demonstrate the versatility of torpor throughout the annual cycle and suggest a fundamental change in physiological feedback between adiposity and torpor during migration. Moreover, this study highlights the underappreciated importance of facultative heterothermy in migratory ecology.

## Introduction

Facultative hypothermia is an energy conservation strategy that allows many mammalian and some avian species to survive periods of resource unavailability or to optimize their energy budgets in certain environments or life stages (McKechnie and Lovegrove, 2002; Ruf and Geiser, 2015). During facultative hypothermia, metabolic rates and body temperatures are reduced to varying extents among species and environmental conditions (Ruf and Geiser, 2015). As some of the smallest avian species, hummingbirds (Trochilidae) are known for their ability to use daily torpor, a deep, short-term form of facultative hypothermia, to cope with energetic challenges they face daily and throughout their annual cycle (Carpenter, 1974; Hainsworth et al., 1977).

Studies that have investigated the use of torpor in hummingbirds in relation to food availability and body mass suggest that torpor initiation is controlled by an endogenous mechanism sensitive to an energy-store threshold (Hainsworth et al., 1977; Hiebert, 1992; Powers et al., 2003). These studies also suggest that the function of torpor shifts seasonally, from an energy emergency survival mechanism to an energy storage maximization strategy during migration (Carpenter and Hixon, 1988; Hiebert, 1993). However, this threshold has not been repeatedly and accurately measured in individual birds spanning life history stages, and the relationship between torpor use and the components of body composition (fat and lean mass) remains unclear.

We explored torpor use in ruby-throated hummingbirds (*Archilochus colubris*), which breed in eastern North America in the early and mid-summer and migrate to wintering grounds in Mexico and Central America in the late-summer. In the breeding period, birds maintain low body masses to optimize aerial agility, important for successful courtship displays and competitive interactions (Altshuler et al., 2010; Hou and Welch, 2016). But like most long-distance migrants, ruby-throated hummingbirds substantially increase their body mass prior to migratory departure to fuel their journey (Hou and Welch, 2016).

We repeatedly and non-invasively quantified the relationship between torpor and endogenous energy stores (fat) in ruby-throated hummingbirds to investigate the underlying rules of torpor use during the breeding and migration periods. We predicted that in the breeding period, torpor initiation would be sensitive to a low energy-store threshold and function as an emergency survival mechanism. We also predicted that this threshold would be abandoned in the late summer to facilitate premigratory fattening. We measured the torpor use and body composition of captive adult (and 1 juvenile) male ruby-throated hummingbirds (n = 16; capture mass: 2.5–3.2 g) that experienced semi-natural photoperiods, on 158 focal bird-nights throughout the summer. We used respirometry to calculate rates of energy expenditure, and quantitative magnetic resonance (QMR) to measure body composition (Guglielmo et al., 2011; Lighton, 2008). We identified the start of torpor entry and arousal by evaluating the slope of each smoothed 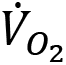 trace, and we calculated the fat content at the time of torpor entry as the amount of evening fat (g) minus the estimated amount of fat expenditure prior to torpor entry, divided by the morning body mass.

Throughout the study period, 13 of 16 birds exhibited low morning body masses until late August and September when they substantially increased their body mass; the birds subsequently maintained high body masses. We analyzed daily changes in morning body mass and defined ‘breeding’, ‘fattening’, and ‘migration’ periods, respectively (Methods, Supplementary Fig. 7). Additionally, 3 ‘non-fattener’ birds maintained a relatively constant body mass throughout the summer (Fig. 1; Supplementary Table 1). QMR scans indicated that changes in body mass were driven primarily by increases in fat r(109)=0.94, 95% CI [0.92, 0.96] P <0.001; Supplementary Fig. 1A), and that body mass and lean mass were slightly negatively correlated (r(107)=− 0.26, 95% CI [−0.43, −0.08], P =0.006; Supplementary Fig. 1B).

**Fig. 1.**
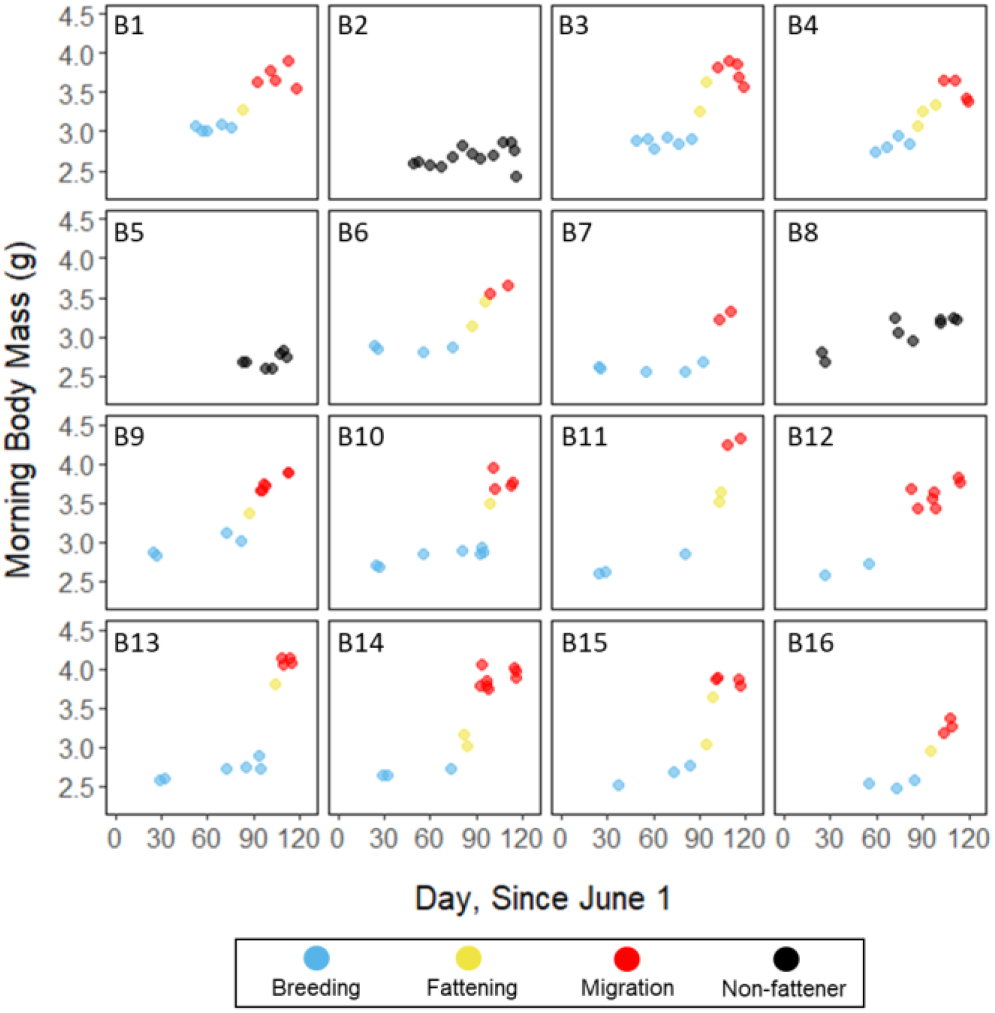
Morning body masses following focal observation nights for each individual bird throughout the study period, with points colored by period.

To determine the rules governing the use of torpor and whether these differed between the breeding, fattening and migration periods, we evaluated changes in the relationships between evening body composition and torpor occurrence, torpor duration, time of torpor entry, fat content at torpor entry, and amount of energy expended before torpor entry, within and among each period. To specifically evaluate the role of torpor in driving premigratory increases in body mass, we compared the magnitude and duration of the fattening period with mean torpor duration and mean daily food consumption during the fattening period.

## Results and Discussion

During the mid-summer breeding period, birds maintained consistently low morning body masses (2.77 ± 0.05 g; slope: 0.001 ± 0.001 g·day^−1^; P=0.34; Fig. 1; Supplementary Table 1). On average, birds used torpor on 61.6 ± 11.1% of focal bird-nights (Supplementary Table 4). When birds started the night with less fat, they were more likely to enter torpor (slope: −0.70 ± 0.22; P=0.001; Supplementary Fig. 2), and they remained torpid longer (slope: −0.66 ± 0.09 hr·%fat^−1^, P<0.001; Fig. 2A). Furthermore, when birds started the night with greater fat content, they entered torpor later in the night (slope: 5.9 ± 0.7 %night·%fat^−1^; P<0.001), expended more energy before initiating torpor (slope: 0.43 ± 0.07 kJ·%fat^−1^, P<0.001, Supplementary Fig. 3A), and lost more fat mass overnight (slope: 1.1 ± 0.2 mg_fat_·%fat^−1^, P<0.001; Fig. 2A).

**Fig 2.**
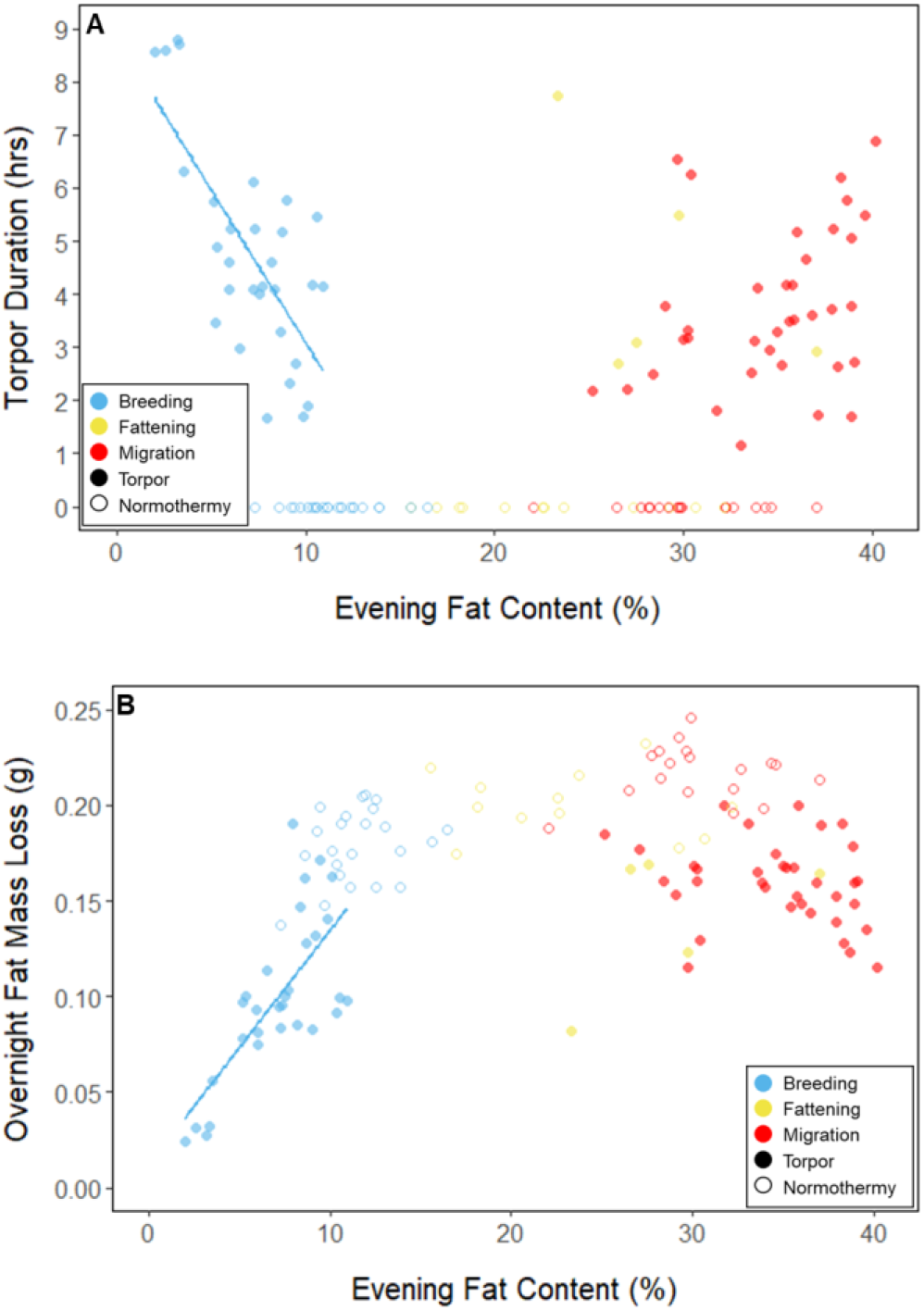
Relationships between evening fat content and (A) torpor duration, and (B) overnight fat mass loss, within each period, with points and significant trendlines colored by period and shaped by torpor use.

Throughout the breeding period, birds consistently entered torpor at a time when they had a relatively low remaining fat level (5.6 ± 0.8 %; Fig. 3B; Supplementary Table 1). Time of torpor entry was not significantly related to the fat level at that time (slope: 0.06 ± 0.03 %fat·%night^−1^; P=0.051; Supplementary Table 4), supporting previous findings that torpor initiation in the breeding season is sensitive to a low constant threshold, irrespective of time of night (Powers et al., 2003) (Fig. 3A). Furthermore, on 47 of 55 (85%) bird-nights, the birds either entered torpor when their fat contents reached approximately 5% of body mass, or did not enter torpor if their fat contents passed this threshold before 75% of the night had elapsed (hummingbirds rarely entered torpor within the last 25% of the night (Hiebert, 1992)) (Supplementary Fig. 6). These results confirm that in the summer breeding period, torpor is sensitive to critically low endogenous energy reserves.

**Fig 3.**
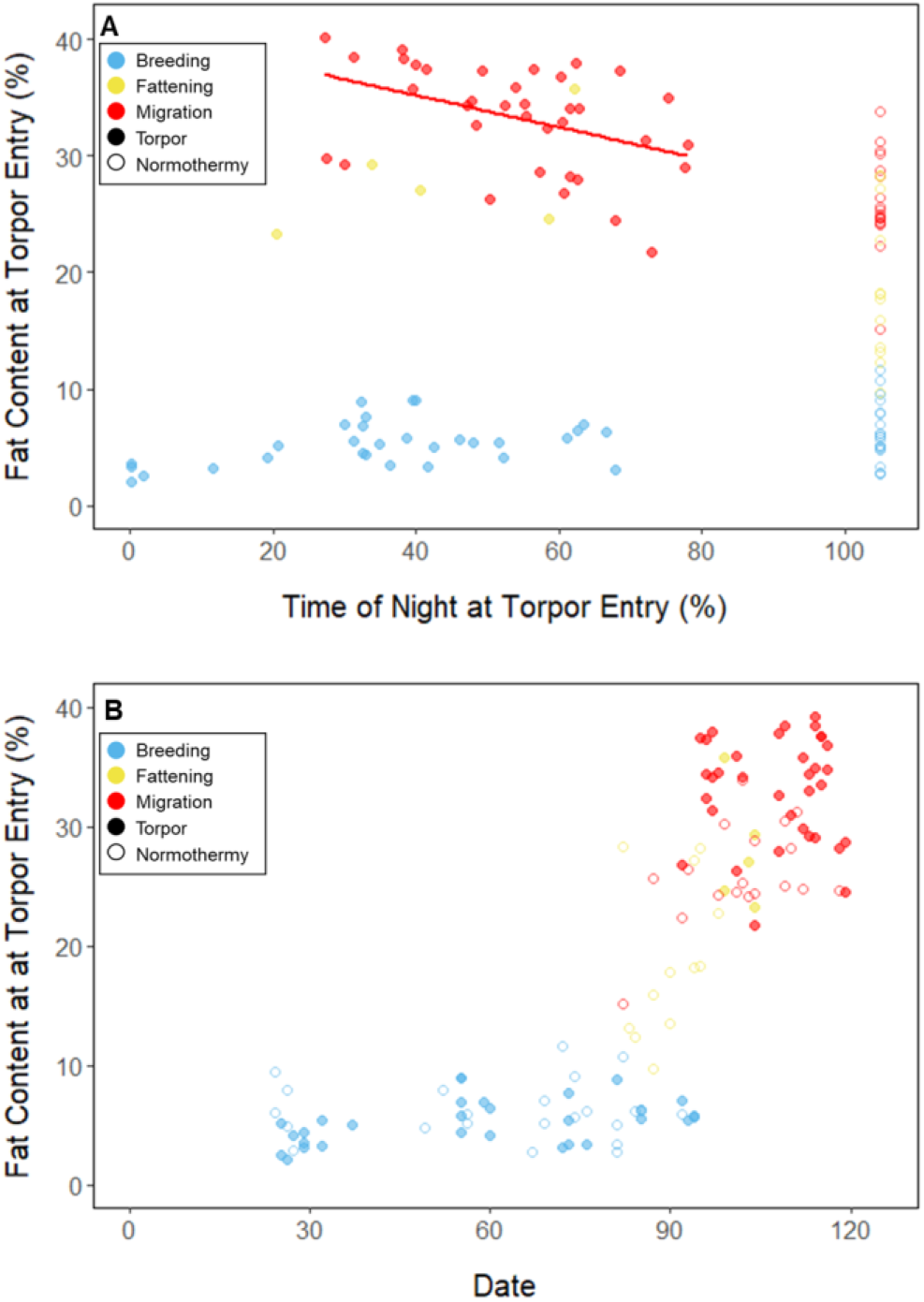
Relationship between fat content at the time of torpor entry (%) and (A) the time of torpor entry (as % of night), and (B) Date, with points and significant trendlines colored by period and shaped by torpor use.

Substantial changes in torpor use accompanied changes in body composition in the late summer when the birds fattened prior to migration. In late August and September, 13 of 16 birds increased body mass (slope: 0.019 ± 0.005 g·day^−1^, P<0.001; Supplementary Table 1) over the span of 10 ± 1 days (range: 6-18 days; Fig. 1; Supplementary Table 3). In this short period, birds increased body mass by an average of 0.58 ± 0.05 g (range: 0.34-0.90 g). Relative to the start of fattening, birds increased their body mass by an average of 19.6 ± 1.6% (range: 11.6-29.0%; Supplementary Table 3).

The magnitude and timing of fattening in these captive birds resembled those documented in free-living ruby-throated hummingbirds (∼0.65g, or 17% over 4 days) (Hou and Welch, 2016). Previous work suggested that both reduced overnight mass loss achieved by torpor use and increased midday foraging effort drive premigratory fattening (Hou and Welch, 2016). To test this hypothesis, we first confirmed the negative relationship between torpor duration and overnight fat mass loss in every period (slopes: breeding: −1.7 ± 0.0 mg_fat_·hr^−1^, fattening: −1.7 ± 0.1 mg_fat_·hr^−1^, migration: −1.5 ± 0.1 mg_fat_·hr^−1^, P<0.001; Supplementary Fig. 4; Supplementary Table 4). Because longer torpor durations invariably spared more fat, we predicted that more frequent and longer torpor use, in addition to higher food consumption during the fattening period, would enhance the rate of premigratory fattening. Most interestingly, birds that used torpor for longer on average achieved greater mass gains during the fattening period (slope: 0.07 ± 0.01 g·hr^−1^; P=0.004; n=11; Fig. 4A; Supplementary Table 2). Contrary to our predictions, food consumption did not significantly affect the magnitude of fattening (slope: 0.00 ± 0.14 g·mL^−1^, P=1.0), and neither torpor duration (slope: −0.6 ± 0.7 hr·days^−1^, P=0.48) nor food consumption (slope: −0.9 ± 0.9 mL·days^−1^, P=0.43) was related to the length of the fattening period (Fig. 4; Supplementary Table 2).

**Fig. 4.**
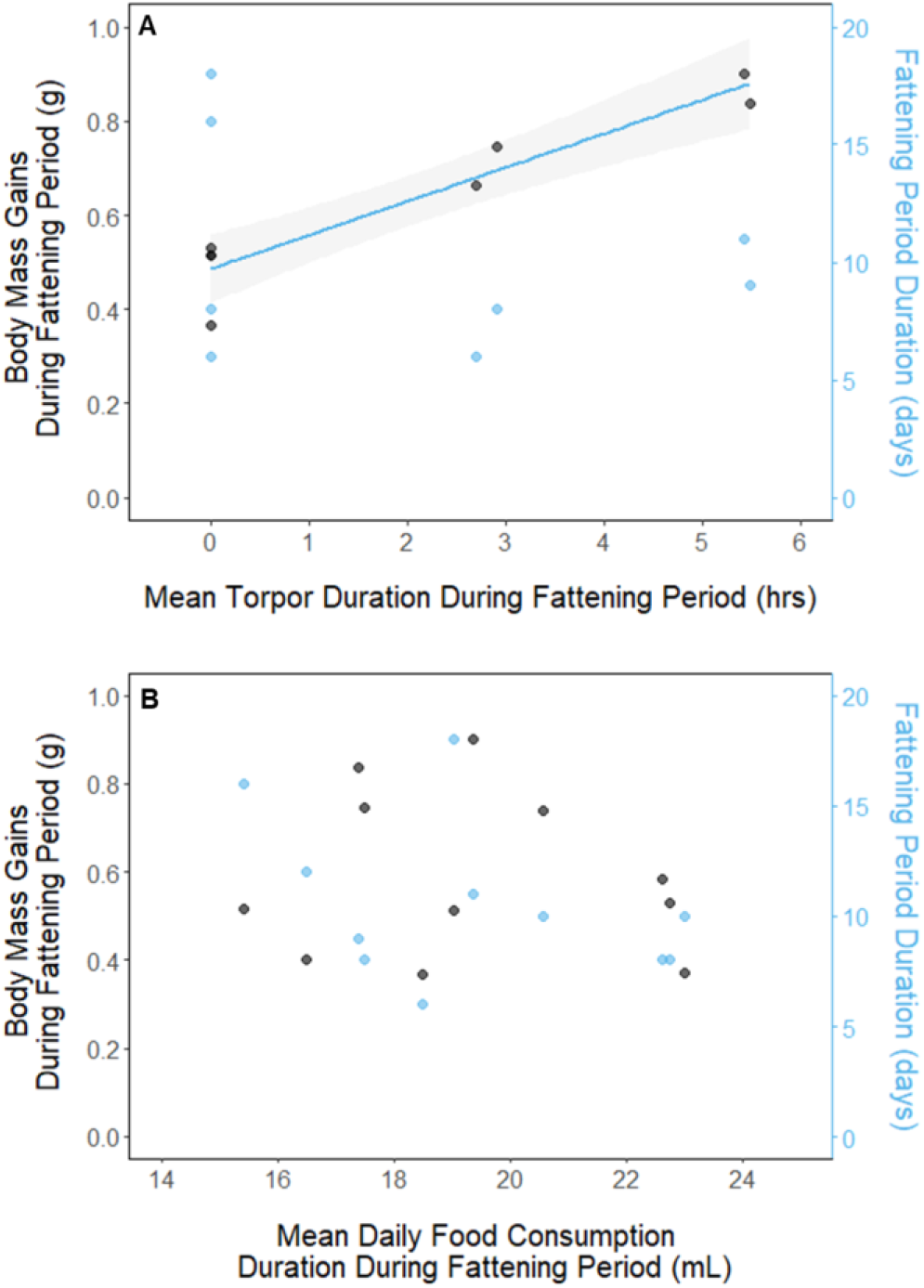
Relationships between magnitude of increases in body mass (black, left axis) and fattening period duration (blue, right axis), and (A) mean torpor duration, and (B) mean daily food consumption, within the fattening period. Trendlines are shown for significant slopes and the shaded area represents 95% confidence intervals.

Birds maintained greater morning body masses in the migration period compared to the summer (3.73 ± 0.05 g, p<0.001), and the migration period body mass remained stable (slope: 0.004 ± 0.002 g·day^−1^, P =0.08; Fig. 1; Supplementary Table 1). Despite beginning nights with 3 to 5 times more fat than they would need to remain normothermic for the entire night at 20°C (0.2 ± 0.01g) and not approaching the breeding period critical threshold, birds used torpor in the migration period at similar frequencies to those of the breeding season (65.1 ± 11.1%, P=0.9; Supplementary Table 4). However, in stark contrast to the breeding period, birds were more likely to enter torpor on nights when they started with greater fat stores (slope: 0.25 ± 0.11; P=0.019; Supplementary Fig. 2). Additionally, the time of torpor entry (slope: −0.8 ± 0.6 %night·%fat^−1^; P=0.20), the amount of energy expended before torpor entry (slope: −0.06 ± 0.05 hr·%fat^−1^; P=0.20; Fig. 3A), torpor duration (slope: 0.10 ± 0.07 hr·%fat^−1^; P=0.13; Fig. 2A), and overnight fat mass loss (slope: 0.2 ± 0.1 mg_fat_·%fat^−1^; P =0.18; Fig. 2B) were not significantly related to evening fat content (Supplementary Table 4).

Fat content at the time of torpor entry was substantially greater in the migration period compared to the breeding period (32.9 ± 0.8%; P <0.001; Fig. 3B; Supplementary Table 4). Additionally, there was a significant negative relationship between time of torpor entry and the fat content at that time (slope: −0.14 ± 0.4 %fat·%night^−1^, P<0.001; Fig. 3A; Supplementary Table 4). While it does not appear that torpor initiation in the migration period is simply sensitive to a high energy threshold, the abandonment of the emergency threshold indicates that the rules governing of torpor use are dependent life history stage, and that hummingbirds may employ torpor as part of various energy management strategies throughout the annual cycle.

In the summer breeding period, ruby-throated hummingbirds reserve torpor for times when they face critically low fat stores during the night. This supports the hypothesis that torpor use is sensitive to a low, consistent threshold of fat in breeding period birds, and that torpor is an energy emergency survival strategy mechanism that protects hummingbirds from depleting energy stores during the night or before they can reach their first meal in the morning (Hainsworth et al., 1977; Hiebert, 1992; Powers et al., 2003). This evidence supports the existence of a ‘adipostat’ mechanism that initiates compensatory physiological changes depending on the level of endogenous energy stores and the amount of fuel demand during a particular season (Boyer and Barnes, 1999; Powers et al., 2003). Although these processes have been well studied in mammals, further avian endocrine studies are needed to elucidate the proximate factors of torpor initiation across taxa. The similarities between the life histories of North American hummingbirds and bats provides an important lens through which to interrogate the ultimate drivers of torpor use throughout the annual cycle (McGuire et al., 2012).

The drastic shift in the relationship between body composition and torpor use, and of the link between torpor duration and mass gains in the late summer compellingly support the ‘torpor assisted migration’ hypothesis, first posited in bats (Carpenter and Hixon, 1988; Hiebert, 1993; Hou and Welch, 2016; McGuire et al., 2012). Unlike most nocturnal migrant birds that can replenish fuel stores during the day, hummingbirds are diurnal migrants that do not forage at night. As in bats, foraging, migrating, and sleeping are mutually exclusive endeavours, so hummingbirds must use a different strategy if they are to reach their destination faster and without making extended refueling stopovers. Thus, using torpor while roosting allows these animals to maintain high fuel stores when feeding opportunities are constrained by migratory priorities.

This is the first study to repeatedly and accurately define a consistent rule governing torpor use in hummingbirds: birds will enter torpor when their fat stores reach a consistently low fat threshold (5%), during life history stages where a lean body composition is advantageous. This rule likely reflects the typically low degree of torpor use in dominant individuals and large species, and the more frequent use of torpor in subordinate individuals and smaller species (Powers et al., 2003). Moreover, our study confirms long standing, but heretofore unproven, assumptions of the key role of torpor in premigratory fattening and in refueling at migratory stopover sites. Hummingbirds dramatically shift their rules of torpor and enter torpor at high fat levels during times when it is advantageous to accumulate excess energy stores. These findings demonstrate the versatility of torpor as an energy management mechanism throughout the annual cycle and have important implications for understanding the physiological basis of torpor initiation.

## Materials and Methods

### Study Animals

Adult (and 1 juvenile) male ruby throated hummingbirds (*Archilochus colubris;* n = 16; capture mass: 2.54–3.2 g) were captured with a modified box trap (drop door trap) in London, ON, Canada, at the University of Western Ontario. Captive hummingbirds were housed individually in EuroCage enclosures (Corners Ltd, Kalamazoo, MI, USA), measuring 91.5 cm W × 53.7 cm H × 50.8 cm D, at the University of Western Ontario’s Advanced Facility for Avian Research. Once captive, birds were fed ad libitum on a 20% (*w*/*v*) solution of a Nektar-Plus (Guenter Enderle, Tarpon Springs, FL, USA). Birds experienced semi-natural photoperiods that were changed approximately weekly, ranging from 15-h light/9-h dark to 12-h light/12-h dark. These photoperiods are reflective of the birds’ natural summer photoperiod, as the data were collected in the summers 2018 and 2019. Lights were abruptly turned on in the morning and shut off in the evening. The birds transitioned from a breeding condition in the beginning of the study period to a migratory condition in end of the study period. Details of animal husbandry and all experiments were approved by the University of Toronto (protocol # 20011649) and the University of Western Ontario Animal Care Committees (protocol #2018-092). Hummingbirds were captured under Ontario Collecting Permit SC-00041.

### Body Composition

This study uses quantitative magnetic resonance (QMR) to measure body composition. QMR is a technology developed in the last 15 years that allows for non-invasive measurement of the masses of fat, lean tissue, and total body water (Guglielmo et al., 2011). QMR allows for short scan times (2-3 minutes), high precision and accuracy, and the ability to measure resting, non-anaesthetized animals (Guglielmo et al., 2011). We used an Echo-MRI (Echo Medical Systems, Houston, TX, USA) with an A10 antenna for measuring birds < 10g. We calibrated the QMR machine with 1.5 g canola oil and 10 g water standards and scanned these standards daily to check the calibration; scans were set at 3 accumulations. On focal nights, birds were scanned 3-5 times in the evening and the morning; the means of the values from these scans were calculated. QMR, paired with respirometry allows us to non-invasively and accurately estimate the level of endogenous energy stores throughout the night, and specifically at the time of torpor initiation.

### Respirometry

This study uses respirometry to calculate rates of energy expenditure. Oxygen consumption and carbon dioxide production rates overnight were obtained via push-flow respirometry using an FC-1B oxygen analyser, a CA-2A carbon dioxide analyser, (Sable Systems International, Las Vegas, NV, USA). Air was first passed through a dew point generator set at 10-15°C and then was flowed into the chambers through Bev-a-line tubing at a rate of 150 mL/min. The excurrent airstream was subsampled at 50 mL/min. Sub-sampled air first passed through a water vapour meter, which measured water vapour pressure (kPa) (RH-300; Sable Systems International). The air then passed through a column containing indicating Drierite (W.A. Hammond DRIERITE, Xenia, OH, USA), the carbon dioxide gas analyser, and the oxygen analyser. Analogue voltage outputs from the thermoresistor, oxygen and carbon dioxide analysers, flow meter, water vapour pressure and in-line barometric pressure sensors were recorded at 1s intervals over the duration of the trial (9-12 h) using EXPEDATA software (v.1.9.27; Sable Systems International) and were recorded on a laptop computer.

Raw data were corrected to standard temperature and pressure, and rates of oxygen consumption 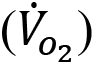 and carbon dioxide production 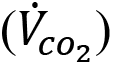 were calculated in Expedata using standard equations (^*12*^; equations 10.6, 10.7, respectively). The rate of oxygen consumption 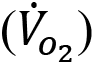, the respiratory exchange ratios (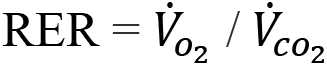, indicates primary metabolic fuel type), and the oxyjoule equivalent were used to calculate the rate of energy expenditure (kJ/min) 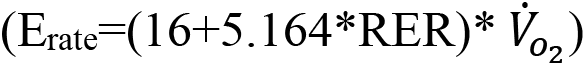 (Lighton, 2008). Where RER was extraneously below 0.71 or above 1.0 it was bound at these limits to satisfy the assumptions of this equation. Total nighttime and metabolic state-specific energy expenditures were calculated by integrating the metabolic rate (*E_rate_*) over time.

### Experimental Protocol

In each overnight experiment, birds were food-deprived 1-3 hours prior to lights-off to ensure crop stores were emptied and the only available sources of energy were endogenous fat and lean mass. Body composition was measured using QMR before the birds were placed in respirometry chambers (10 cm W × 10 cm H × 20 cm D) at 20°C and the lights were turned off. Torpid birds could not be scanned because disturbing them would cause them to unnaturally arouse and because their cooler body temperatures could decrease scan precision and consistency (Guglielmo et al., 2011). Immediately following lights-on in the morning, body composition was measured. Air temperature was measured directly outside the chamber via a thermoresistor; although air temperature inside the chamber was not recorded, experiments show the inside and outside air temperatures were within 1°C.

### Data Processing

Between June and September 2018 and 2019, we recorded repeated overnight measurements on 15 adult male birds and 1 male juvenile, for a total of 158 bird nights. The birds began exhibiting increased body masses (indicative of migratory condition) at different times in the late summer (mean: August 28; range: August 12- September 8). In order to identify periods of distinct rates of change in body mass, we analyzed the rate of change in morning body mass across the study period (Supplementary Fig. 7). We first smoothed the morning body mass trace with a smoothing parameter (0.35) that we identified through an iterative process. We calculated the first derivative of this smoothed body mass trace, and defined periods based on bird-specific ‘cut-off’ slopes that we calculated as 75% of the maximum slope (g/day). We defined ‘breeding’ as periods where the rate of change was less than the cut-off slope, ‘fattening’ as periods where the rate of change was greater than the cut-off slope, and ‘migration’ as periods where the rate of change was less than the cut-off slope and after the start of the fattening period. When this analysis yielded periods that clearly disagreed with visual inspection of the curve, we slightly adjusted the bounds of the fattening period to fit a more realistic pattern. Three birds that did not fatten were defined as ‘non-fatteners’ and were excluded from the statistical models that regard seasonal changes in torpor use.

We calculated torpor propensity as the percentage of nights the birds entered torpor of the total number of nights for each bird within each period. In order to calculate the energetic characteristics of torpor, such as the fat content at the time of torpor entry and duration, the temporal characteristics of torpor must be clearly defined. In much of the avian and mammalian torpor literature, torpor entry is defined by phrases such as, when the metabolic rate “abruptly declines”, or by criteria such as a threshold value of a set proportion of the average normothermic resting values of body temperature or metabolic rate (Ruf and Geiser, 2015; Shankar et al., 2020; Wolf et al., 2020). However, these various and often vague definitions are problematic for repeatability and our understanding of energy metabolism at specific stages of torpor. In order to identify accurate and repeatable periods of consistent metabolic states, we analyzed the rate of change in 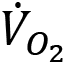 across the night (Supplementary Fig. 8). We first the linearly interpolated 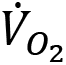 and smoothed this trace using a smoothing parameter (0.6) that we identified through an iterative process. We calculated separate smoothed traces for the time before (containing entry) and after (containing arousal) the end of torpor/start of arousal. To determine this intermediate point we calculated the derivative of a smoothed trace of the entire night (using night-specific smoothing factor), and preliminarily identified the approximate end of torpor/ start of arousal as the minute the rate of change was greater than an arousal cut-off slope of 0.005 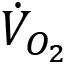 /min. We then calculated the first derivative of each of the entry and arousal smoothed 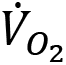 traces, and defined states based on entry and arousal cut-off slopes. We defined ‘entry’ as periods where the rate of change was greater than two standard deviations of the rate of change during the normothermic period before torpor (which was identified by analyzing the preliminary smoothed curve where the rate of change was less than 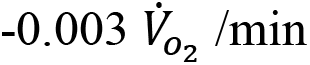). We defined ‘arousal’ as periods where the rate of change of was greater than four standard deviations higher than the mean rate of change during the normothermic period before torpor; we defined the end of the arousal period as the point when the bird exhibited peak 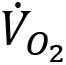. We defined ‘torpor’ as periods where the 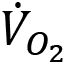 was stable and between entry and arousal. We also annotated points before the start of entry and after the end of arousal as ‘normo-pre’ and ‘normo-post', respectively. This process yielded accurate and repeatable metabolic state annotations of each minute.

The initial evening and the final morning percent fat content were calculated as fat mass (g) / body mass (g). We used the rate of energy expenditure, cumulative energy expenditure and initial fat mass measurements to estimate instantaneous percent fat content throughout each night. These data allowed for the estimation of the amount of energy reserves, relative to body mass, of each bird at the time of torpor initiation. We calculated this value by subtracting the fat mass equivalent (1g_fat_/37kJ) of cumulative energy expenditure at torpor entry from evening fat mass and dividing the result by morning body mass. We calculated torpor duration (h) as the time between the start of torpor entry until the beginning of arousal (excluding arousal). We calculated overnight mass losses as the fat mass equivalent of the amount of overnight energy expenditure. Lastly, we calculated body mass increases as the change in body mass from the beginning to the end of the fattening period, relative to the smoothed morning body mass trace used to determine the periods.

## Statistical Analyses

We used mixed effects analyses to evaluate the relationships between various response and predictor variables, while accounting for intra-individual differences (Bates et al., 2015). First, we used linear mixed effects analyses to determine how body mass, fat content, lean mass, and food consumption changed within and among (*with respect to* date) each period of distinct mass change (with day length as a covariate for the food consumption model). We evaluated the overall relationships between body mass and fast mass, and body mass and lean mass, using a repeated measures correlation test (‘*rmcorr*’ R package; (Bakdash and Marusich, 2017)).

We used a linear mixed effects model to compare mean torpor propensity among periods, and a logistic mixed effects model to compare the influence of evening fat content on probability of occurrence of torpor within each period. We also used linear mixed effects models to determine how torpor duration, energy expenditure before torpor entry, and overnight fat mass loss changed with respect to evening fat content within and among periods, with night duration as a covariate. We used linear mixed effects models to determine how % fat at the time of torpor entry changed with respect to date and percent of night duration within and among periods. We also used linear mixed effects models to determine how overnight mass loss changed with respect to torpor duration within and among periods. Lastly, we used linear models to evaluate the effect of mean torpor duration and mean daily food consumption within the fattening period on the magnitude and duration of the fattening period.

We performed all statistical analyses using R (R Core Team, 2020). To generate linear and logistic mixed effects models, we used the ‘*lme4*’ package, with Bird ID as a random effect (Bates et al., 2015). For linear models we used the ‘*lm*’ function. For each response variable, we iteratively compared several combinations of relevant fixed effects, and used the AICs to determine the most parsimonious model. The model with the lowest AIC by at least 2 points was considered the best, and we verified that residuals of this model showed homoscedasticity and normality. We determined *P*-values using the ‘*anova*’ function on each model generated using the ‘*lmerTest*’ package, and made pair-wise comparisons within and among periods using the ‘*emmeans’* package. All values are given as estimated marginal means ± standard error, unless otherwise indicated, and significance was taken at α < 0.05.

## Acknowledgments

We thank AFAR staff (Francis Boon and Michela Rebuli), and undergraduate assistants. We also thank Welch lab members (University of Toronto), Morag Dick, and Anusha Shankar for discussions. We also thank Alex Gerson and his lab members for providing housing, and for discussions. This work was supported by a Natural Sciences and Engineering Research Council of Canada (NSERC) Discovery Grant 06129-2015 RGPIN and Human Frontier Science Program (number RGP0062/2016) to KCW; and NSERC Discovery Grant 05245-2015 RGPIN, and Canada Foundation for Innovation, Ontario Research Fund to CGG.

## Author contributions

Correspondence and requests for materials should be addressed to EE. E.E., and K.W., contributed to project conceptualization and experimental design. K.W. and C.G. provided resources and funding and advised on technical approach. E.E. performed data collection, curation, analysis, and visualization. E.E. prepared original drafts, which K.W. and C.G. reviewed and edited. All authors approved the final manuscript.

## Supplementary Information

is available for this paper. This contains datasets, R code, and a metadata file.

## Supplementary Figures and Tables

This paper contains Supplementary Figures 1–8 and Tables 1–4

## Data and materials availability

All data is available in the main text or the supplementary materials. Analyses reported in this article can be reproduced using the data and code provided by Eberts et al., (2021). (Data, code, and metadata file will be uploaded to a publicly available data repository after submission).

## Supplementary Figures

**Supplementary Fig. 1.**
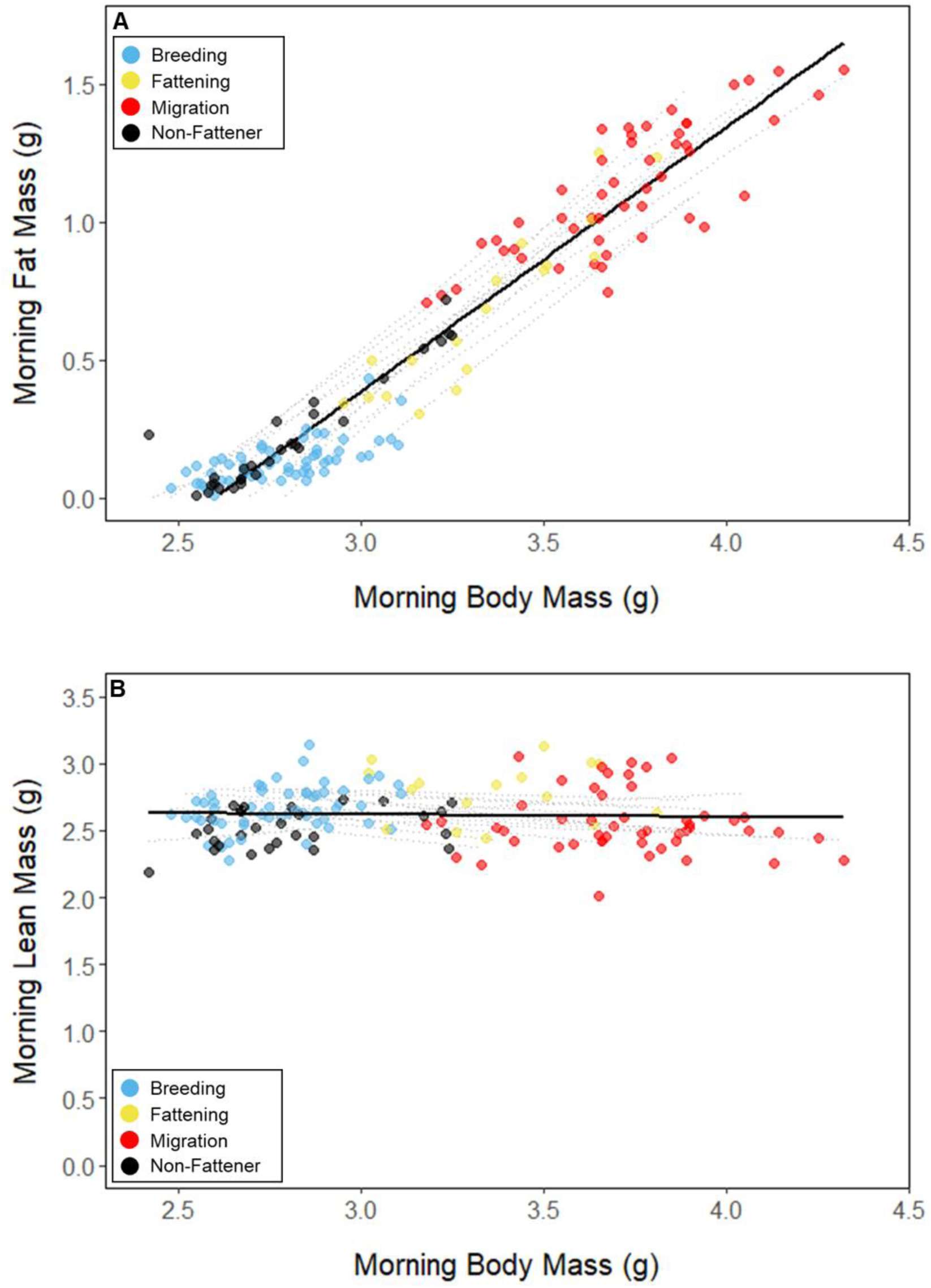
Relationships between body mass and (A) fat mass, and (B) lean mass, with points colored by period. Black lines represent the overall linear relationship during the entire study period and dashed lines represent bird specific regressions.

**Supplementary Fig. 2.**
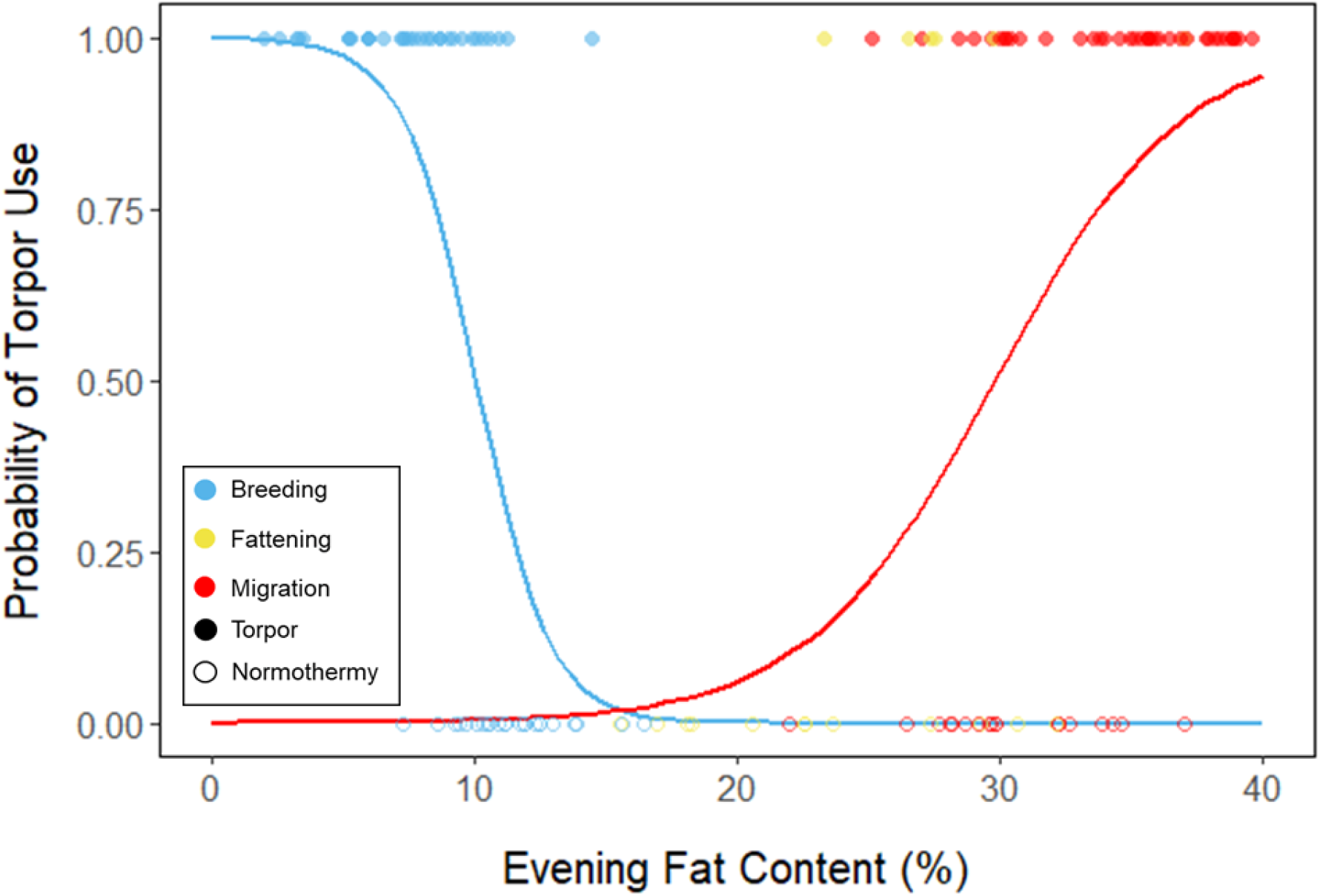
Logistic regression of torpor use as a function of evening fat content, with points and significant trendlines colored by period and shaped by torpor use. Lines are predicted logistic curves estimating the probability of torpor use with respect to evening fat mass.

**Supplementary Fig. 3.**
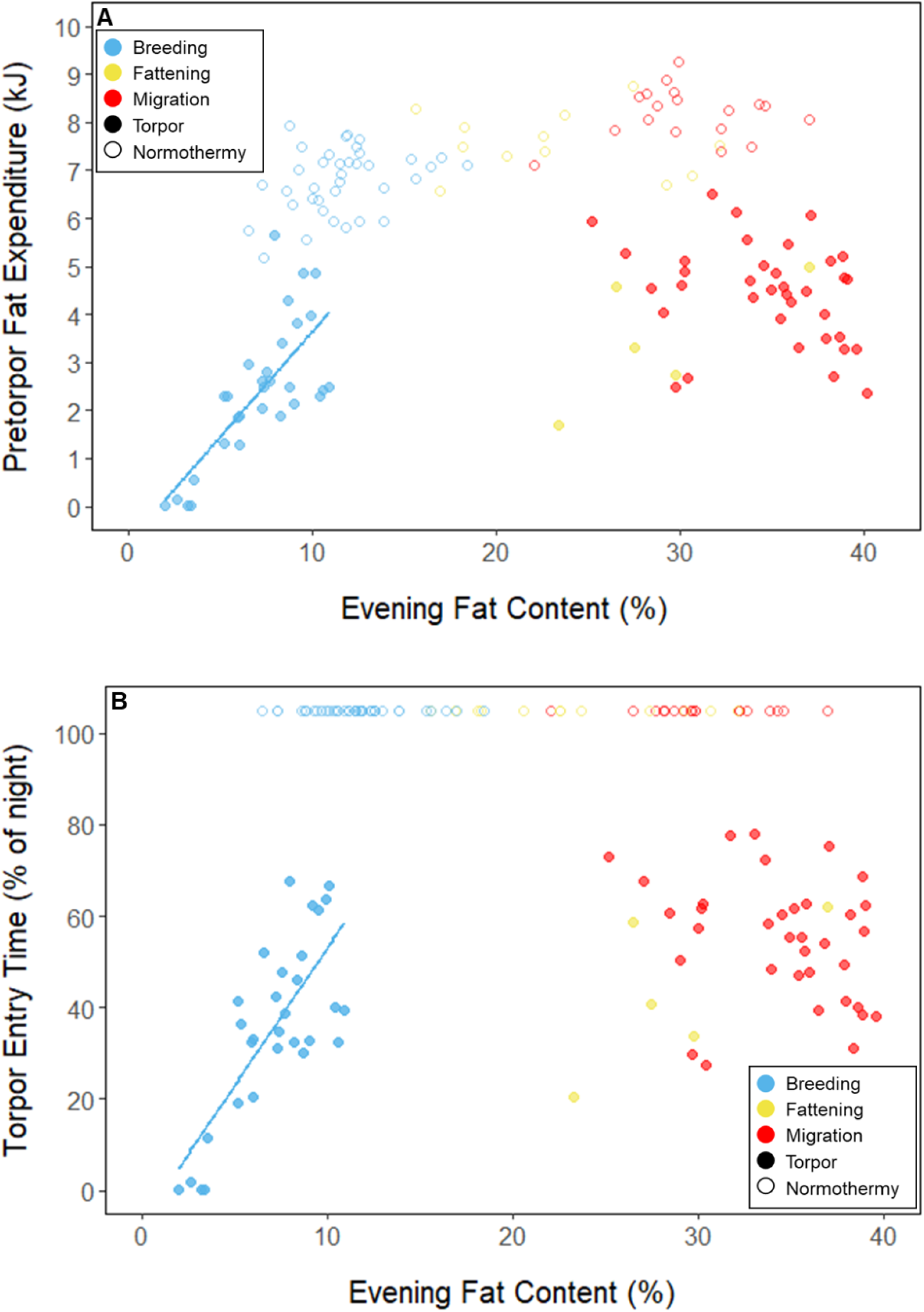
Relationships between evening fat content and (A) energy expenditure before torpor entry, and (B) time of torpor entry, within each period, with points and significant trendlines colored by period and shaped by torpor use.

**Supplementary Fig. 4.**
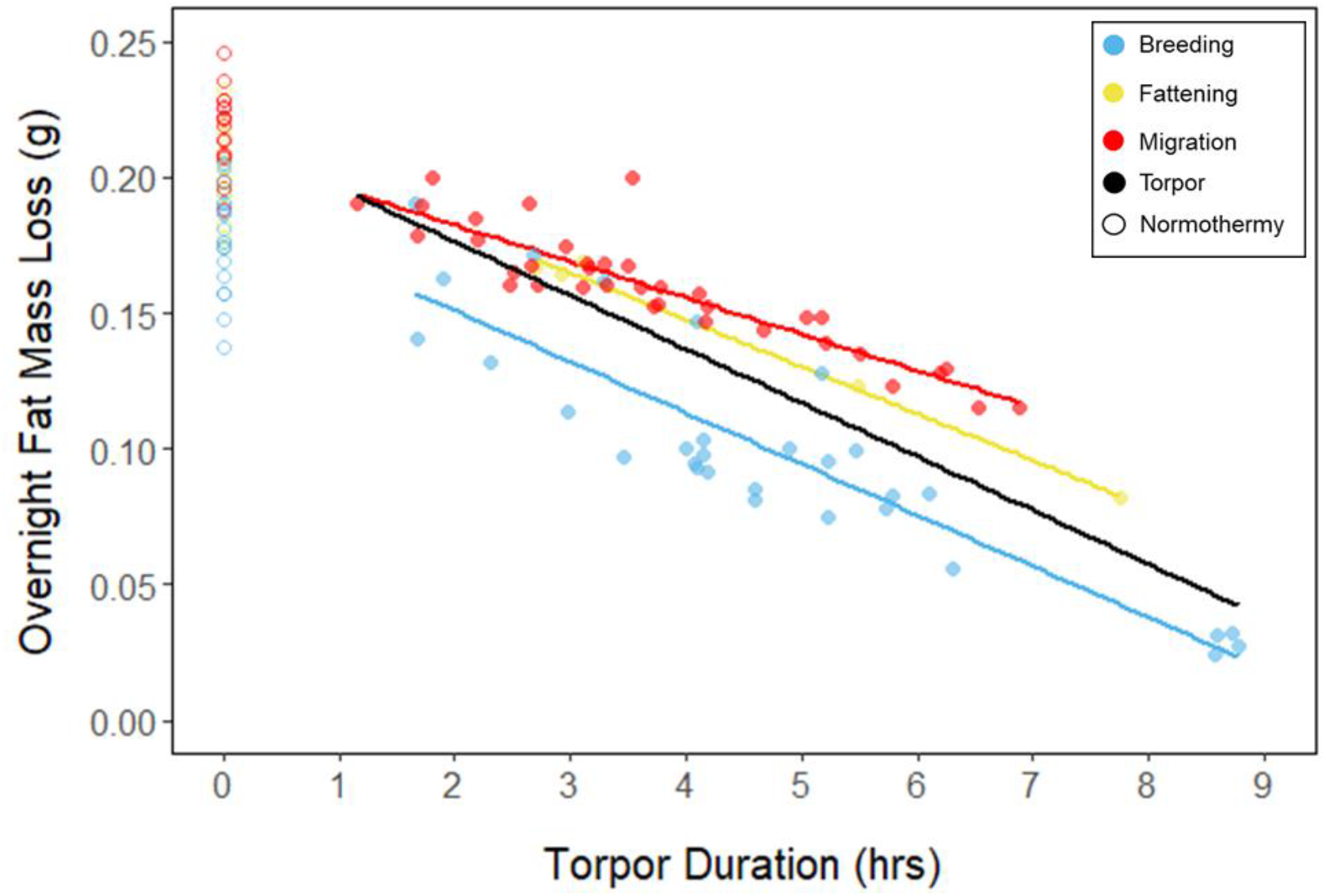
Relationships between torpor duration and overnight fat mass loss, within each period, with points and significant trendlines colored by period and shaped by torpor use. Black line indicates overall linear relationship irrespective of period.

**Supplementary Fig. 5.**
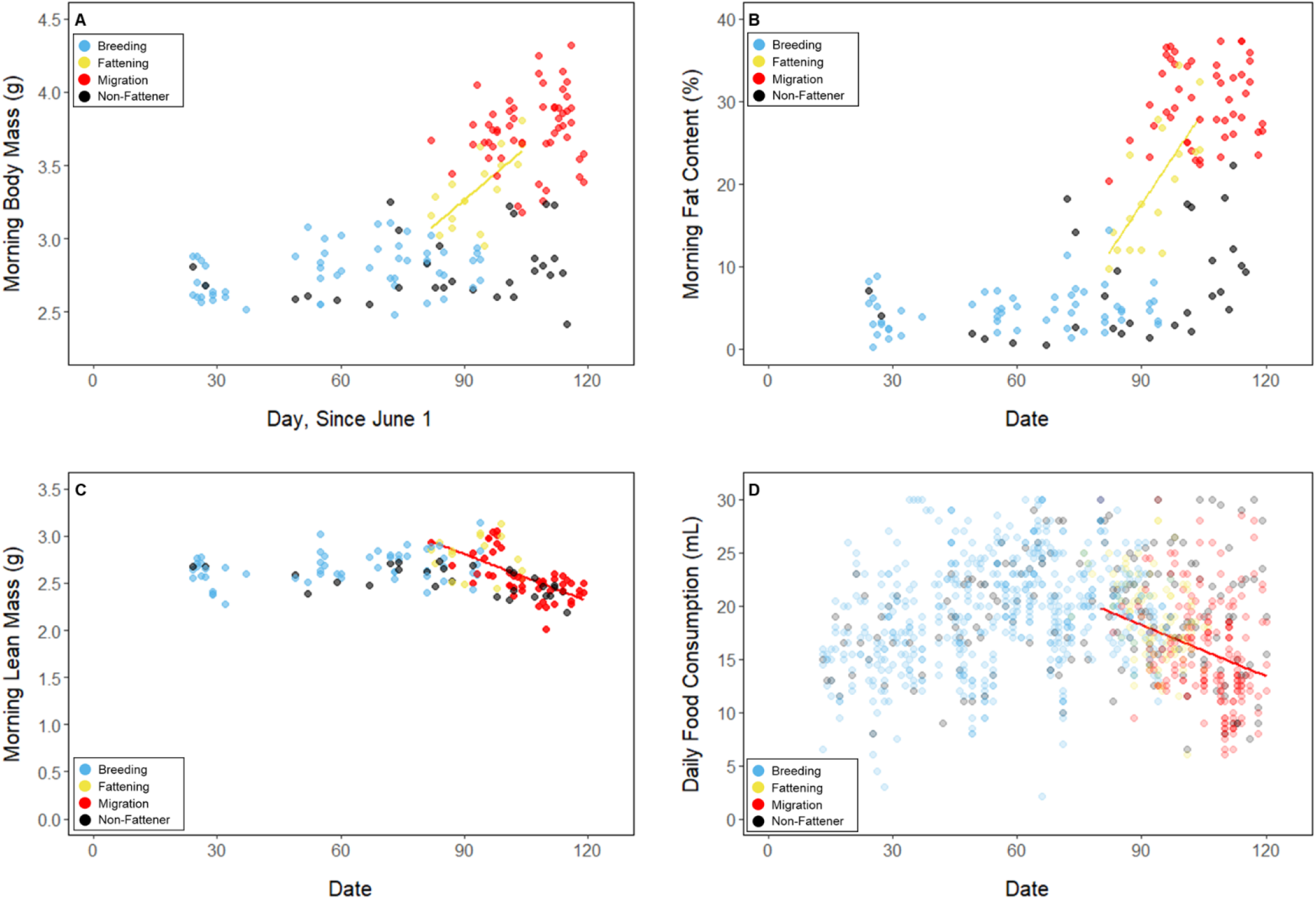
(A) Body mass, (B) fat content, (C)lean mass, on the mornings following focal observation nights, and (D) daily food consumption throughout the study period, with points and significant trendlines colored by period.

**Supplementary Fig. 6.**
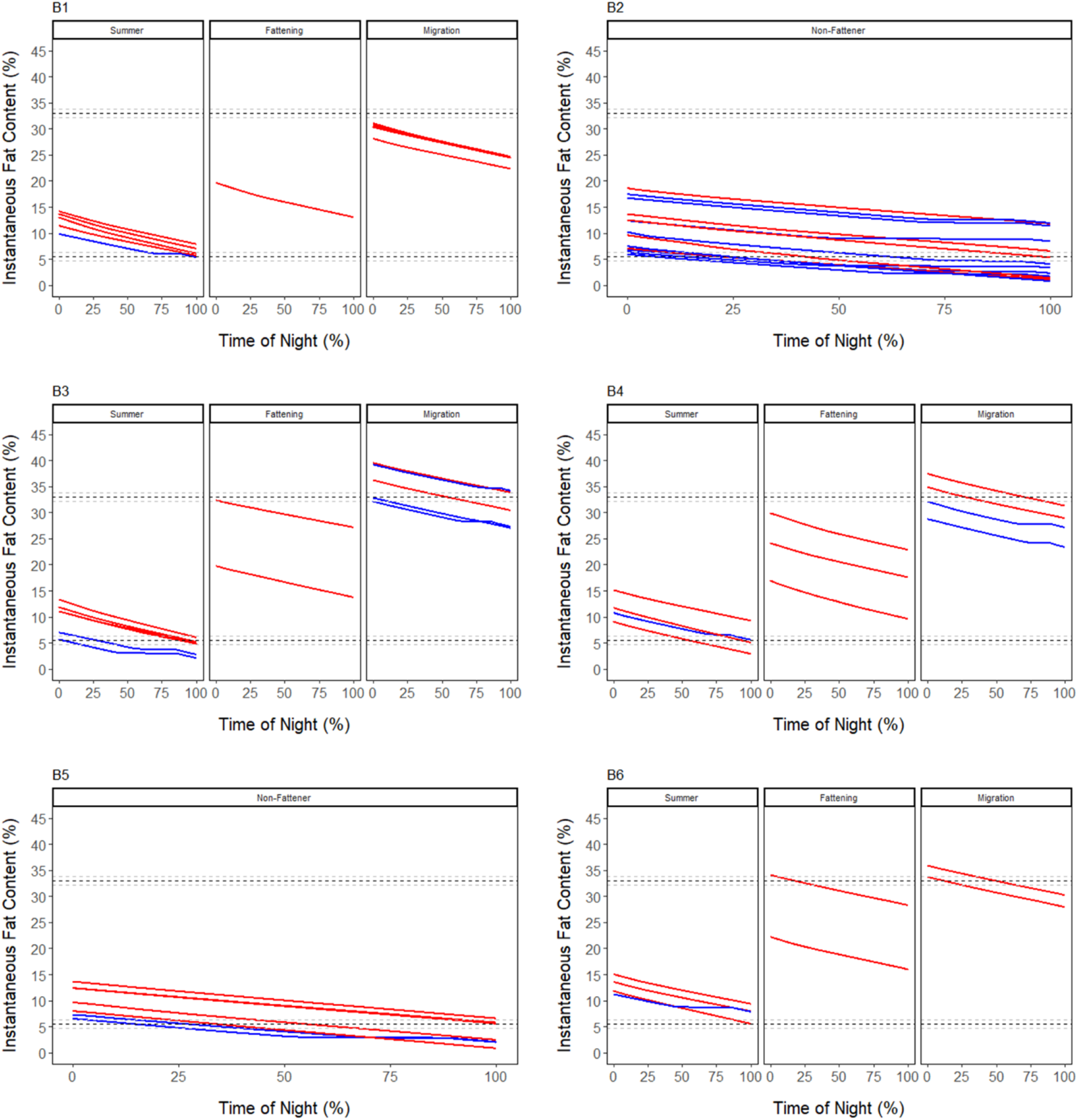
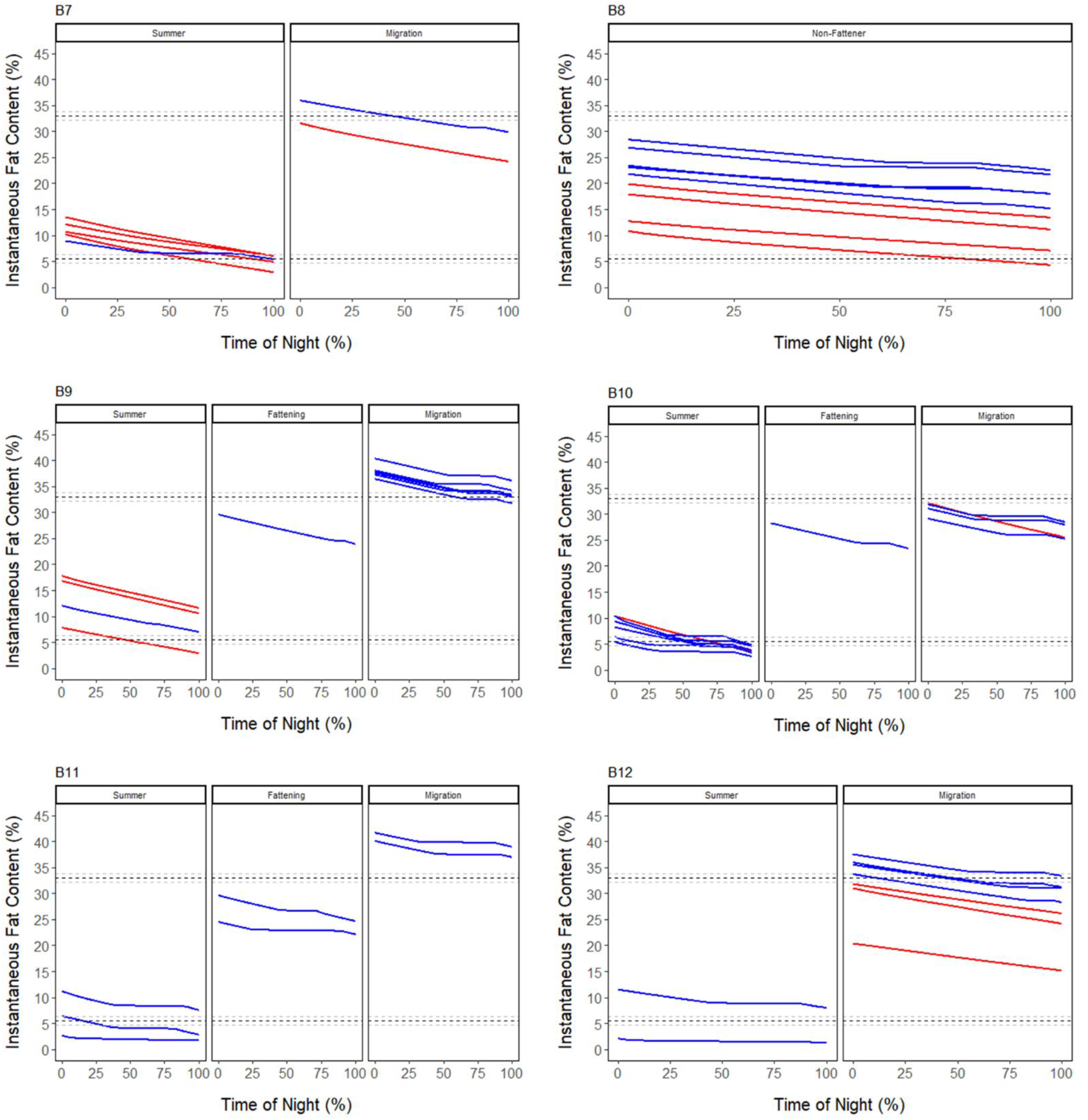
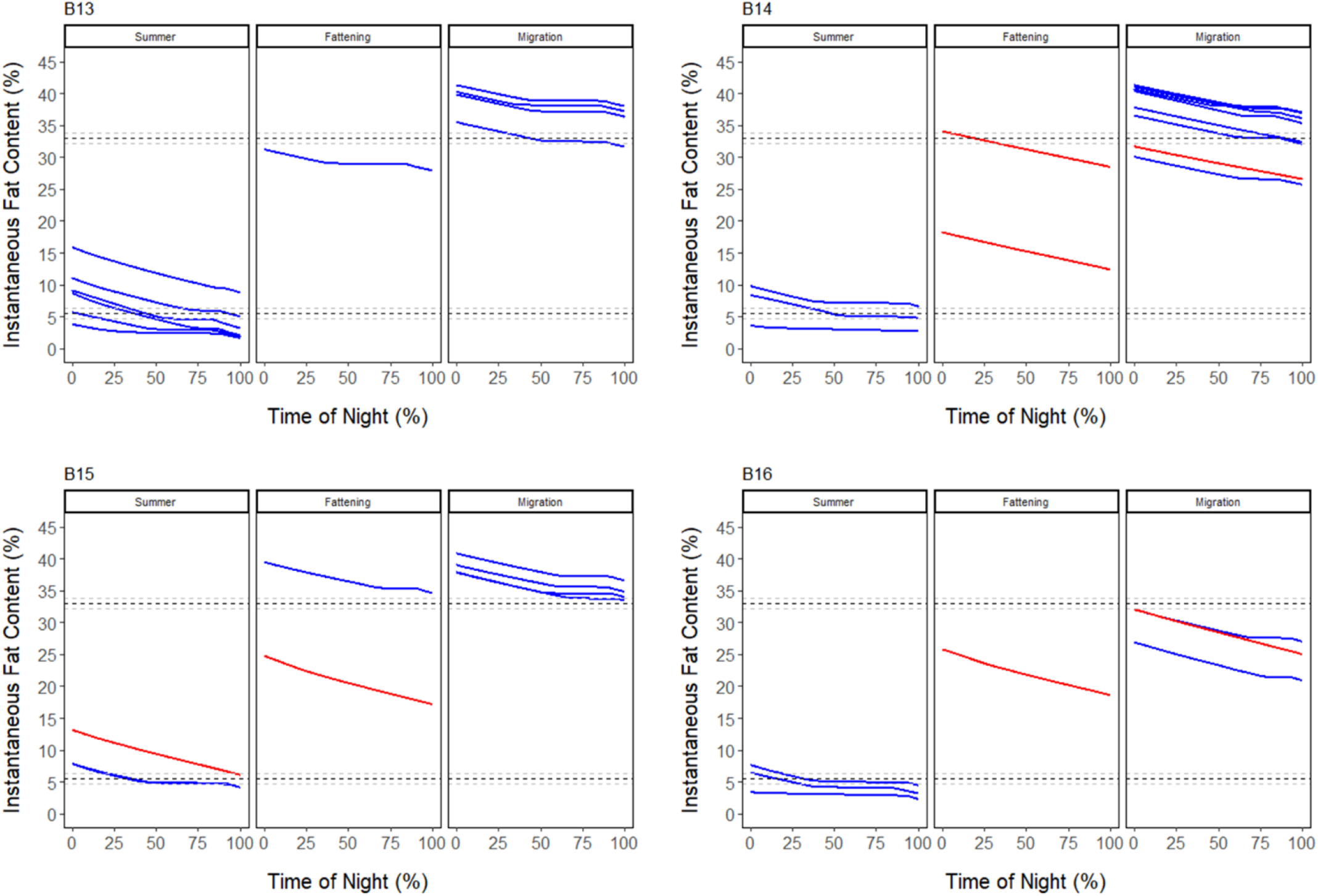
Instantaneous percent fat content over time (as % of night) throughout each summer, fattening, and migration period, as well as during the whole study period for non-fatteners. Red lines represent normothermic nights and blue lines represent torpid nights. The average breeding threshold ± 1 standard error is indicted by horizontal dashed black and grey lines, respectively.

**Supplementary Fig. 7.**
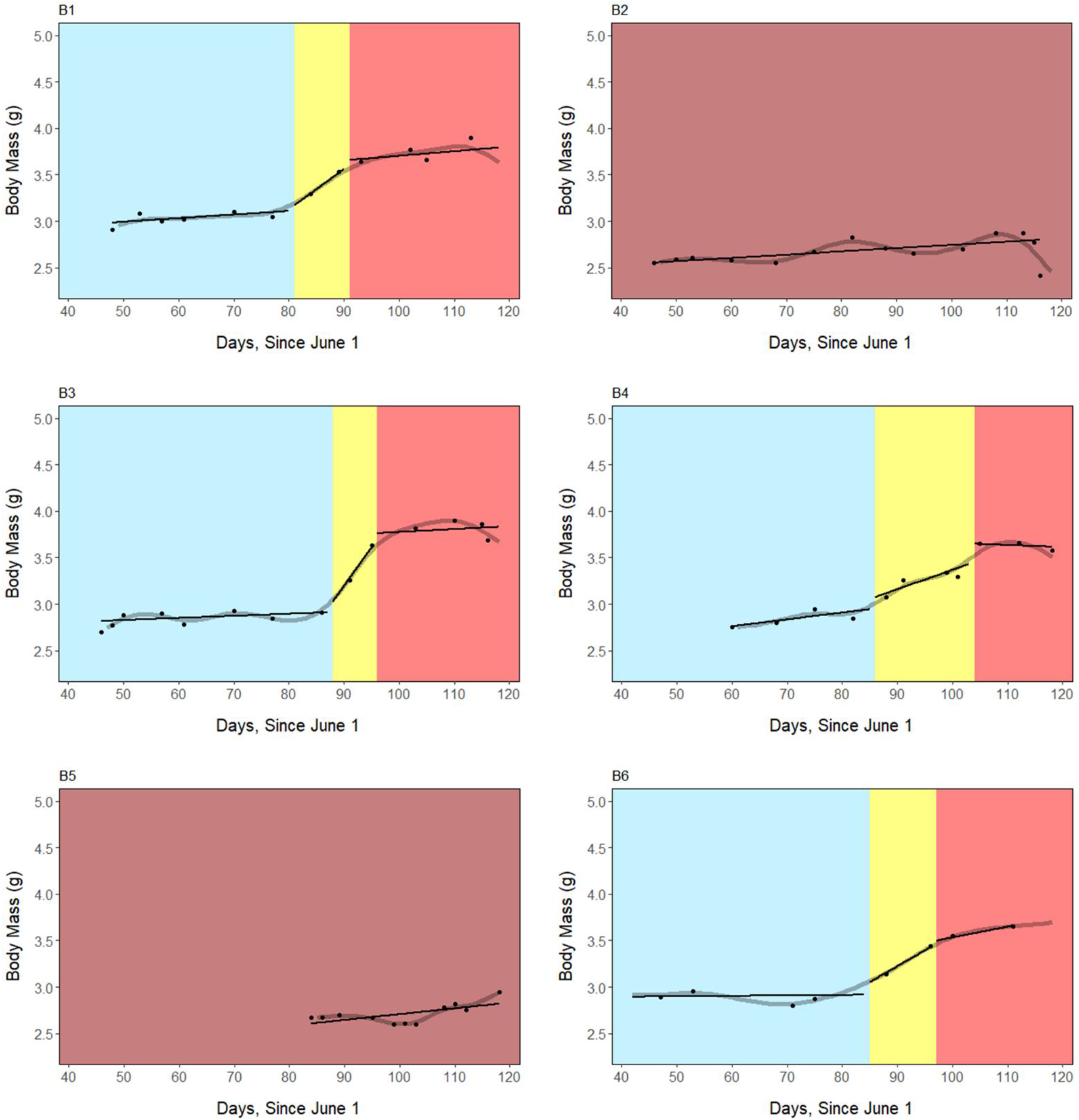
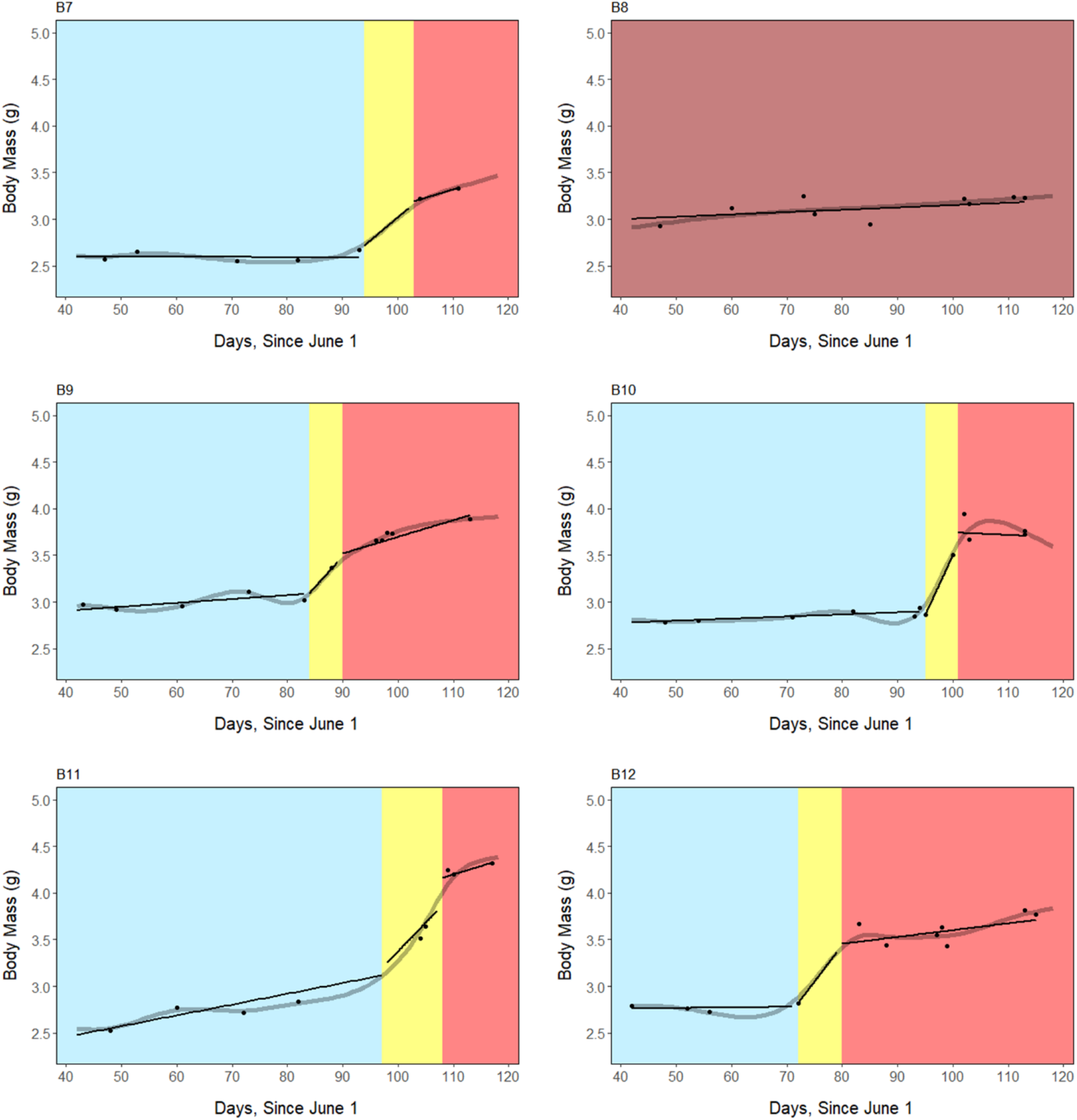
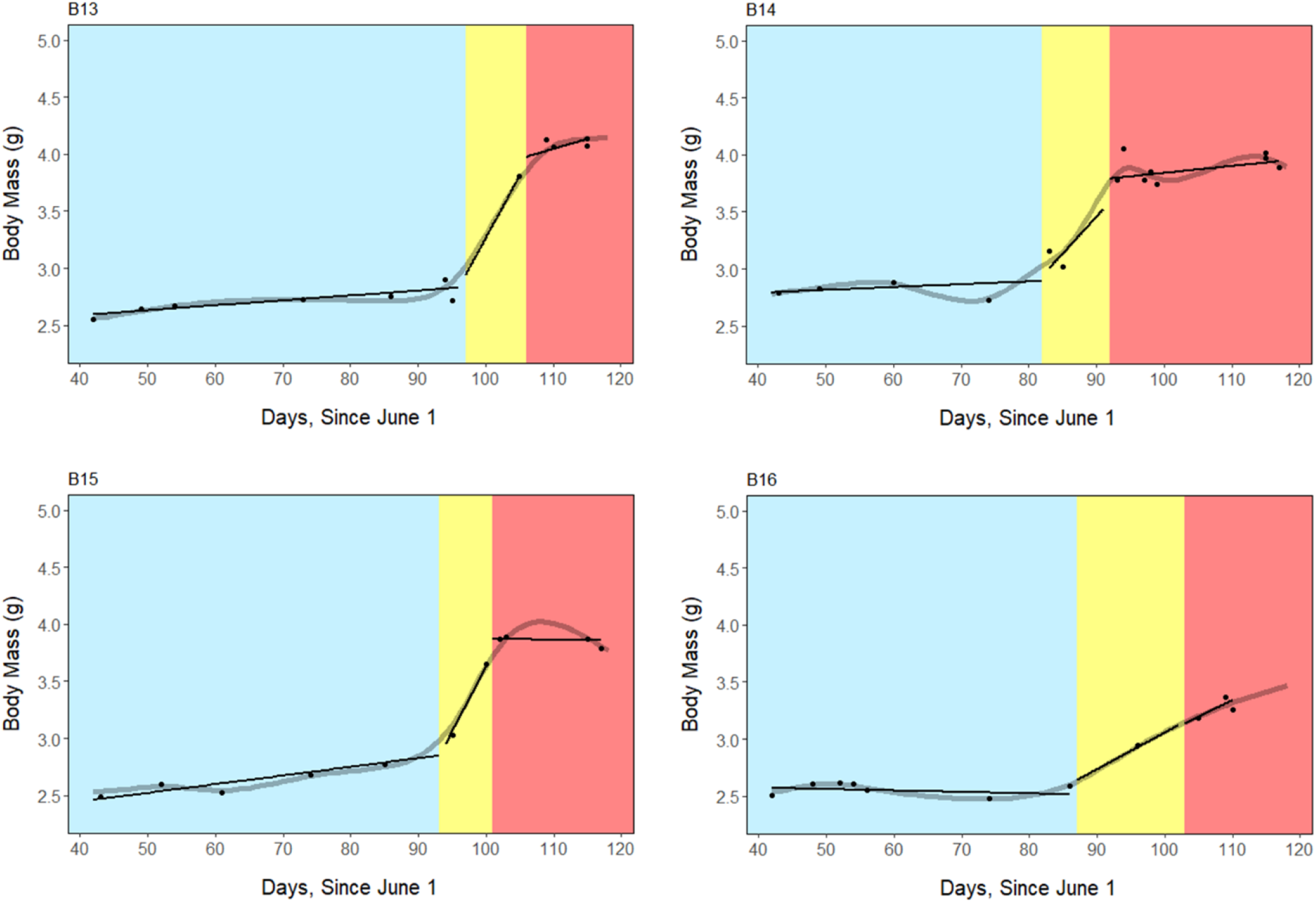
Morning body mass (black points) following focal nights across the entire study period, starting at the date of capture. These data points were smoothed (grey line), and the slope of these points was used to define breeding, fattening, and migration periods for each bird, which are shaded blue, yellow, and red, respectively. Non-fatteners are also included and shaded dark red.

**Supplementary Fig. 8.**
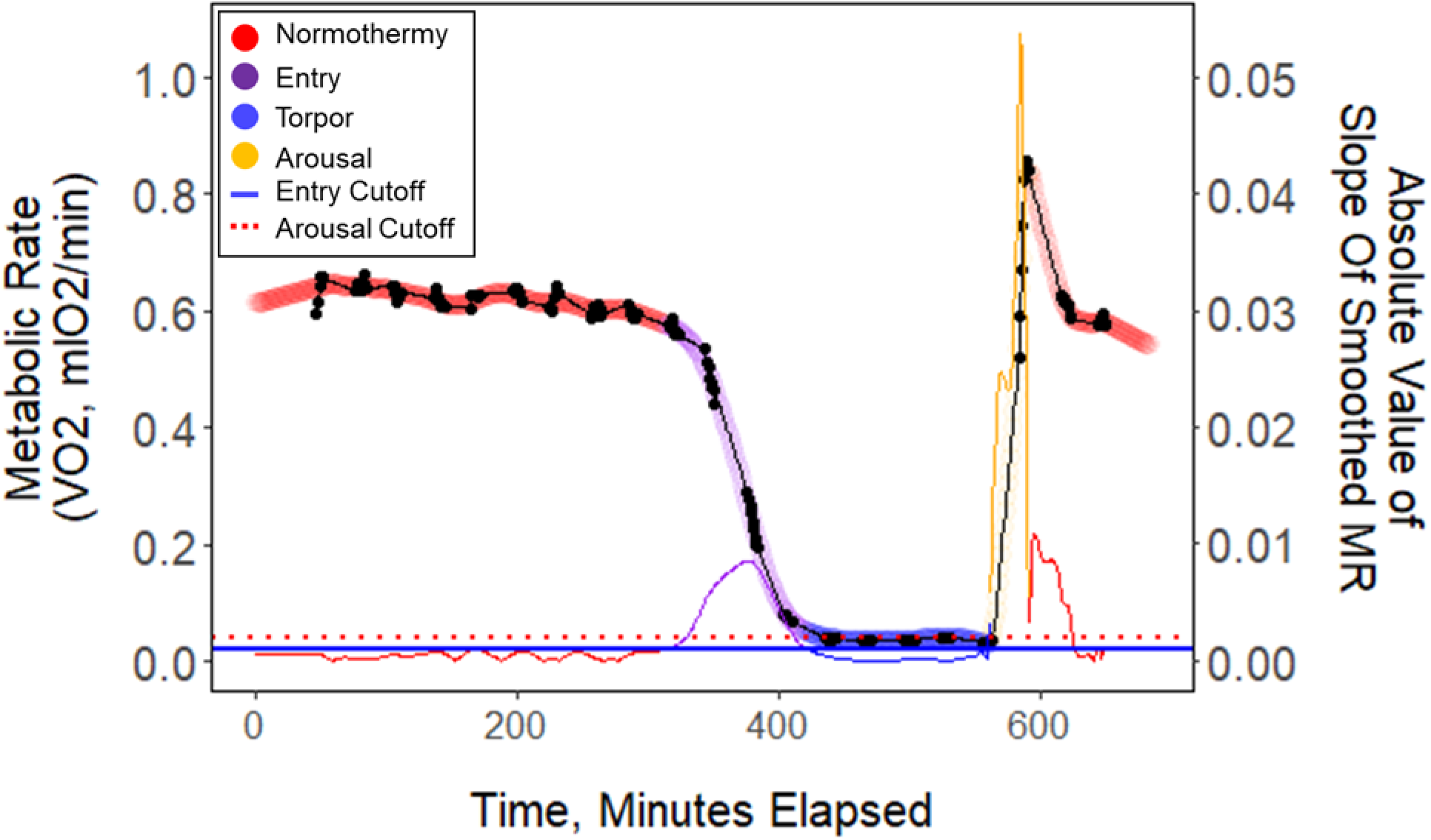
Raw 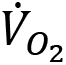 data (black points) plotted every minute throughout one exemplar night of torpor (B13_9.2.19). These data points were smoothed (thick colored line), and the slope of these points (thin colored line) was used to determine metabolic state. Horizontal lines represent entry (blue) and arousal (red) cut-off slopes which were used to distinguish metabolic states of each minute.

## Supplementary Tables

**Supplementary Table 1.** Estimated marginal means and slopes of body composition and food consumption variables, descriptions of the statistical models, and statistical comparisons within and among periods.

| Model <sup>a</sup> | Body Mass |  |  | Fat Content (% fat mass/body mass) |  |  |
| --- | --- | --- | --- | --- | --- | --- |
|  | ~ Period | ~ Period *<br>Date |  | ~ Period | ~ Period *<br>Date |  |
| Period (n) | Mean ± SE<br>(g) | Slope ± SE<br>(g/day) | *p* | Mean ± SE<br>(%) | Slope ± SE<br>(%/day) | *p* |
| Breeding<br>(13) | 2.77±0.05<br>(F, M) <sup>b</sup> | 0.001±0.001<br>(F) | 0.34 | 4.7±0.8<br>(F, M) | 0.04±0.02<br>(F) | 0.053 |
| Fattening<br>(13) | 3.31± 0.06<br>(B, M) | 0.019 ± 0.005* <sup>c</sup><br>(B, M) | <0.001 | 19.6±1.1<br>(B, M) | 0.81± 0.12*<br>(B, M) | <0.001 |
| Migration<br>(13) | 3.73± 0.05<br>(B, M) | 0.004±0.002<br>(F) | 0.08 | 29.9±0.8<br>(B, F) | 0.11±0.06<br>(F) | 0.06 |
| Non-fatteners<br>(3) | 2.81±0.10 | 0.004±0.001 | <0.001 | 7.6±2.7 | 0.15±0.02 | <0.001 |

| Model <sup>a</sup> | Lean Mass |  |  | Food Consumption |  |  |
| --- | --- | --- | --- | --- | --- | --- |
|  | ~ Period | ~ Period *<br>Date |  | ~ Period +<br>Day Length | ~ Period *<br>Date +<br>Day Length |  |
| Period (n) | Mean ± SE<br>(g) | Slope ± SE<br>(g/day) | *p* | Mean ± SE<br>(mL/day) | Slope ± SE<br>(mL/day) | *p* |
| Breeding<br>(13) | 2.67±0.03<br>(M) | 0.002 ± 0.001*<br>(M, N) | 0.02 | 19.8 ± 0.6<br>(M) | 0.95±0.2*<br>(M) | <0.001 |
| Fattening<br>(13) | 2.80±0.05<br>(M) | -0.001±0.006 | 0.85 | 19.4±0.8<br>(M) | 0.4±0.5 | 0.47 |
| Migration<br>(13) | 2.55±0.03<br>(B, F) | -0.016 ± 0.003*<br>(B, N) | <0.001 | 17.0±0.8<br>(B, F) | -0.4±0.4<br>(B) | 0.17 |
| Non-fatteners<br>(3) | 2.52±0.06 | -0.002±0.001 | 0.009 | 20.4±2.3 | 0.5±0.1*<br>Daylength:<br>*p*<0.001 | <0.68 |

<sup>a</sup> Asterix in model indicates interaction.
<sup>b</sup> Statistically significant comparisons within parentheses, alpha taken at 0.05 (P<0.05).
<sup>c</sup> Asterix after value indicates significance.

**Supplementary Table 2.** Absolute and relative increases in body mass, fat mass, and fat content during the fattening period.

| Value | Absolute |  | Relative (%) |  |
| --- | --- | --- | --- | --- |
| | Mean ± SE | Range | Mean ± SE | Range |
| Fat Duration (days) | 10±1 | 6-18 | ----- | ----- |
| Body Mass Increase (g) | 0.58±0.05 | 0.34-0.90 | 19.6±1.6 | 11.6, 29.0 |
| Fat Mass Increase (g) | 0.60±0.05 | 0.34-0.84 | 216.2±28.8 | 83.1, 486.1 |
| Fat Content Increase (%) | 15.0±0.9 | 8.8-19.1 | 163.7±23.4 | 63.7, 390.1 |

**Supplementary Table 3.** Correlations between body mass gains and duration of the fattening period, with mean torpor duration in the fattening period and torpor propensity in the fattening period.

| Model | Body Mass Gains (g) |  | Fat Duration (days) |  |
| --- | --- | --- | --- | --- |
|  | Body Mass Gains (g) ~<br>Mean Torpor Duration (h) +<br>Mean Food Intake |  | Fat Duration (days) ~<br>Mean Torpor Duration (h) +<br>Mean Food Intake |  |
| | Slope ± SE | *p* | Slope ± SE | *p* |
| Mean Torpor Duration (h) | 0.07±0.01*<sup>a</sup> | 0.004 | -0.6±0.7 | 0.48 |
| Mean Weekly Food Intake (mL/day) | 0.00±0.14 | 1.0 | -0.9±0.9 | 0.43 |

<sup>a</sup> Asterix after value indicates significance.

**Supplementary Table 4.**
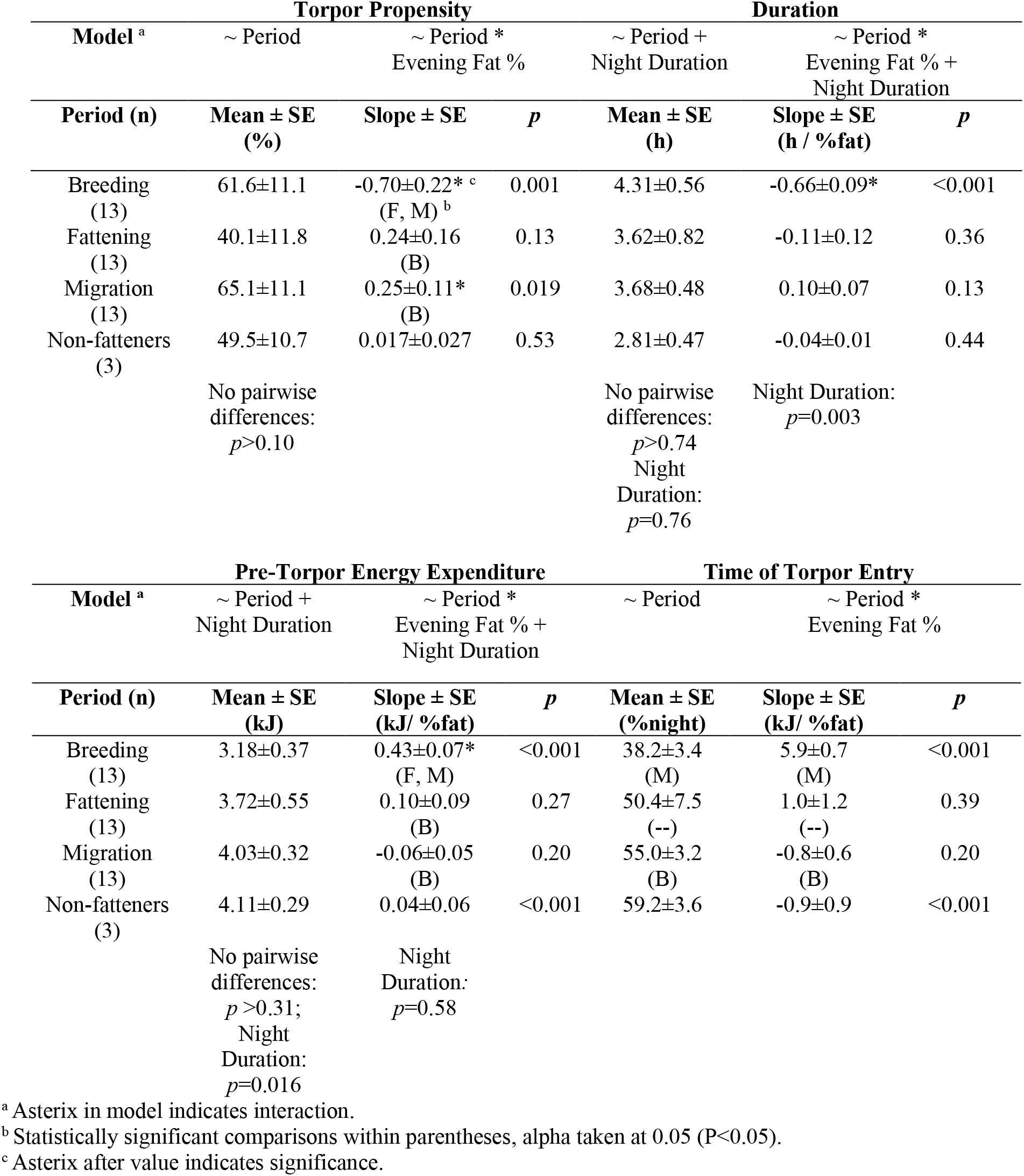
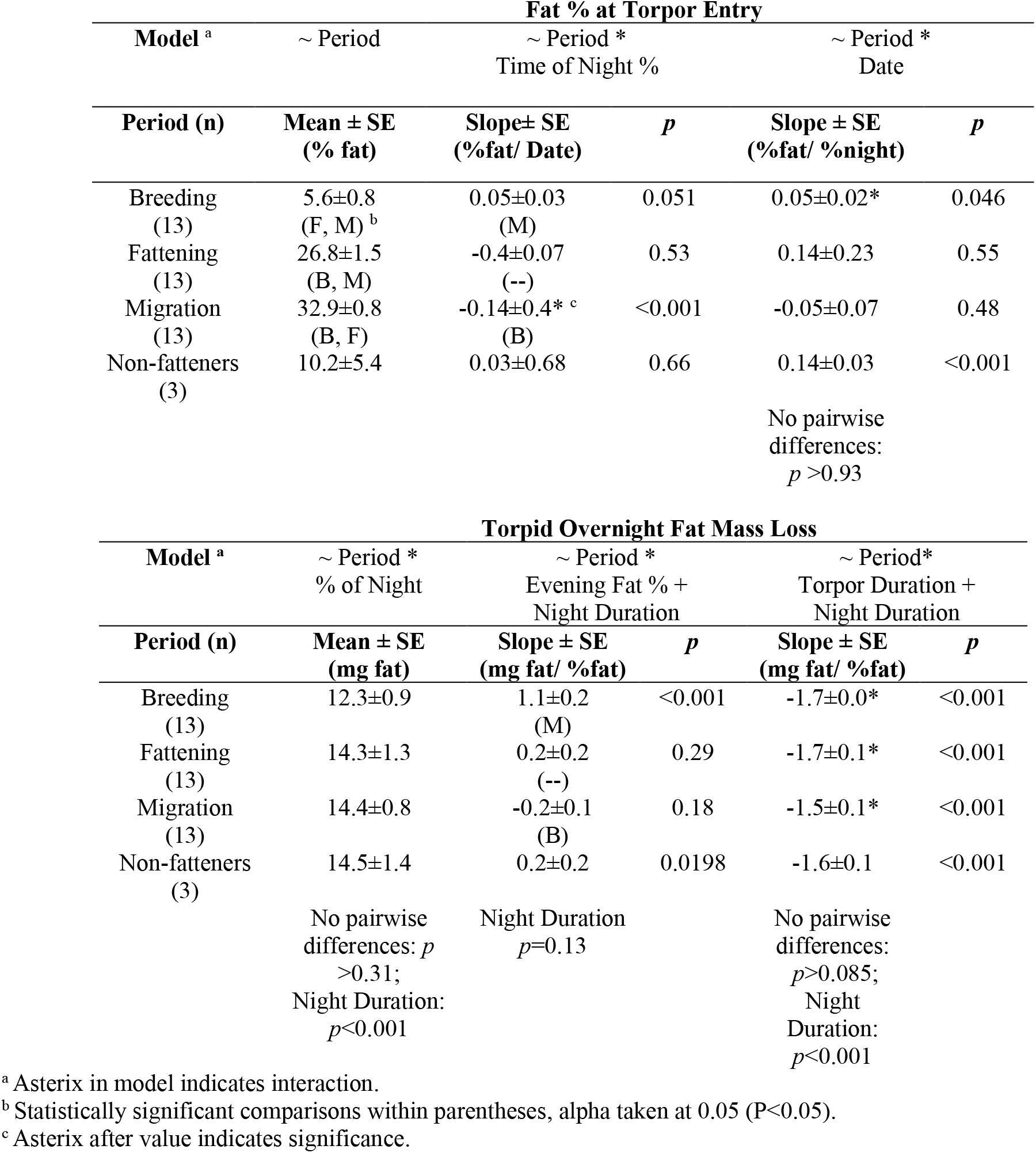
Estimated marginal means and slopes of torpor use variables, descriptions of the statistical models, and statistical comparisons within and among periods.

## References

Altshuler DL, Dudley R, Heredia SM, McGuire JA. 2010. Allometry of hummingbird lifting performance. J Exp Biol 213:725–734. doi:10.1242/jeb.037002

Bakdash JZ, Marusich LR. 2017. Repeated measures correlation. Front Psychol 8.

Bates D, Mächler M, Bolker B, Walker S. 2015. Fitting Linear Mixed-Effects Models Using lme4. J Stat Softw 67.

Boyer BB, Barnes BM. 1999. Molecular and Metabolic Aspects of Mammalian Hibernation. Bioscience 49:713–724. doi:10.2307/1313595

Carpenter FL. 1974. Torpor in an Andean hummingbird: Its ecological significance. Science (80−) 183:545–547. doi:10.1126/science.183.4124.545

Carpenter FL, Hixon MA. 1988. A New Function for Torpor: Fat Conservation in a Wild Migrant Hummingbird. Condor 90:373–378. doi:10.2307/1368565

Guglielmo CG, McGuire LP, Gerson AR, Seewagen CL. 2011. Simple, rapid, and non-invasive measurement of fat, lean, and total water masses of live birds using quantitative magnetic resonance. J Ornithol 152:75. doi:10.1007/s10336-011-0724-z

Hainsworth FR, Collins BG, Wolf LL. 1977. The Function of Torpor in Hummingbirds. Physiol Zool 50:215–222. doi:10.1086/physzool.50.3.30155724

Hiebert SM. 1993. Seasonal changes in body mass and use of torpor in a migratory hummingbird. Auk 787–797.

Hiebert SM. 1992. Time-dependent thresholds for torpor initiation in the rufous hummingbird (Selasphorus rufus). J Comp Physiol B 162:249–255. doi:10.1007/BF00357531

Hou L, Welch KC. 2016. Premigratory ruby-throated hummingbirds, Archilochus colubris, exhibit multiple strategies for fuelling migration. Anim Behav 121:87–99. doi:https://doi.org/10.1016/j.anbehav.2016.08.019

Lighton JRB. 2008. Measuring metabolic rates: a manual for scientists. Oxford University Press.

McGuire LP, Guglielmo CG, Mackenzie SA, Taylor PD. 2012. Migratory stopover in the long-distance migrant silver-haired bat, Lasionycteris noctivagans. J Anim Ecol 81:377–385. doi:10.1111/j.1365-2656.2011.01912.x

McKechnie AE, Lovegrove BG. 2002. Avian Facultative Hypothermic Responses: A Review. Condor 104:705–724. doi:10.1093/condor/104.4.705

Powers DR, Brown AR, Van Hook JA. 2003. Influence of Normal Daytime Fat Deposition on Laboratory Measurements of Torpor Use in Territorial versus Nonterritorial Hummingbirds. Physiol Biochem Zool 76:389–397. doi:10.1086/374286

R Core Team. 2020. R: A Language and Environment for Statistical Computing.

Ruf T, Geiser F. 2015. Daily torpor and hibernation in birds and mammals. Biol Rev Camb Philos Soc 90:891–926. doi:10.1111/brv.12137

Shankar A, Schroeder RJ, Wethington SM, Graham CH, Powers DR. 2020. Hummingbird torpor in context: duration, more than temperature, is the key to nighttime energy savings. J Avian Biol 51:1–14. doi:10.1111/jav.02305

Wolf BO, Mckechnie AE, Schmitt CJ, Czenze ZJ, Johnson AB, Witt CC, Wolf BO. 2020. Extreme and variable torpor among high-elevation Andean hummingbird species 5–9.

